# Optimization of mRNA synthesis and its cell expression for vaccine development

**DOI:** 10.1101/2025.05.13.653856

**Authors:** Subrata K Das, Gaurav Dutt, Vishakha Goswami, Abul Faiz, Alpana Joshi

## Abstract

In vitro transcription (IVT) is a versatile procedure that facilitates template-directed synthesis of RNA molecules of any sequence, from short oligonucleotides to those of several kilobases, in quantities ranging from micrograms to milligrams. This technique, involving the engineering of a template with a bacteriophage promoter sequence, enables the synthesis of RNA for use in a variety of applications including structural studies, biochemical assays, and as functional molecules. Messenger RNA (mRNA) therapy has tremendous potential in regenerative medicine, disease treatment, and vaccination. Synthetic mRNA leverages the cell’s natural translation machinery to produce proteins, with the ability to transfect cells and induce expression of target proteins under physiological conditions until it is eventually degraded. In this study, we explore the impact of pseudouridine (Ψ)-modified mRNA in enhancing RNA stability, translational efficiency, and immune compatibility for therapeutic use. By optimizing IVT conditions and employing cellulose-based purification techniques, we successfully synthesized high-quality, modified mRNA that demonstrated superior functional performance. Luciferase and GFP mRNA, synthesized with pseudouridine-modified rNTPs, exhibited improved stability, reduced immune activation, and enhanced translation efficiency in HEK293 cells. A sevenfold increase in luciferase activity and elevated GFP fluorescence confirmed the higher protein expression capabilities of the modified mRNA. Furthermore, cellulose bead purification effectively separated single-stranded RNA from double-stranded RNA contaminants, ensuring minimal immune response and maximizing transfection efficiency. These findings highlight the potential of pseudouridine-modified mRNA and refined purification methods for advancing mRNA-based therapies, from vaccines to protein-replacement treatments, setting the foundation for scalable, clinical-grade RNA production.

## INTRODUCTION

Messenger RNA (mRNA) has emerged as a pivotal tool in the development of modern therapeutics, offering a transient, non-integrative approach to protein expression. Since its discovery in 1961, mRNA research has significantly advanced, overcoming challenges related to stability, immunogenicity, and delivery [1] [2]. Early studies revealed that direct injection of in vitro transcribed (IVT) mRNA into animal models could result in protein expression, laying the foundation for mRNA-based therapeutic strategies [3]. Unlike DNA-based approaches, which require nuclear entry and pose risks of genomic integration, mRNA functions solely in the cytoplasm, reducing the risk of insertional mutagenesis while enabling rapid and transient protein production [1]. mRNA-based therapeutics have garnered significant attention in the fields of infectious diseases, oncology, and protein-replacement therapies. For example, recent breakthroughs in mRNA vaccine development against COVID-19 showcased the immense potential of mRNA technology [4] [5]. These vaccines utilize chemically modified mRNA to encode viral antigens, prompting robust immune responses. Beyond vaccines, mRNA is being explored for its ability to encode functional proteins, such as enzymes, antibodies, or growth factors, in diverse therapeutic context [6] [7]. However, successful mRNA therapy hinges on several critical factors: efficient in vitro transcription (IVT), effective purification to remove dsRNA contaminants, and reliable delivery systems to ensure cellular uptake and protein translation [8] [9] [10].

The introduction of chemically modified nucleotides, such as pseudouridine (Ψ), represents a transformative milestone in mRNA technology [11] [12]. These modifications improve the stability and translational efficiency of mRNA while mitigating its immunogenicity. Natural RNA, particularly bacterial and mitochondrial RNA, tends to activate innate immune responses through Toll-like receptors (TLRs) such as TLR3, TLR7, and TLR8 [13] [14]. This immune activation, mediated by unmodified RNA, is often associated with the presence of double-stranded RNA (dsRNA) contaminants. Conversely, the incorporation of pseudouridine into RNA reduces its recognition by immune sensors, enabling its therapeutic application without eliciting excessive inflammatory responses [2] [11].

In vitro transcription (IVT) serves as the cornerstone of mRNA production, enabling the synthesis of high-quality RNA templates for therapeutic and research purposes. Key components of the IVT reaction include a DNA template with a T7 promoter, ribonucleotide triphosphates (rNTPs), and T7 RNA polymerase. The addition of pseudouridine-modified rNTPs further enhances RNA stability and translation [15]. Nevertheless, IVT reactions can yield dsRNA byproducts, which are potent activators of the innate immune system through TLR3. Therefore, robust purification protocols are indispensable to separate single-stranded RNA (ssRNA) from dsRNA. Methods such as cellulose-based purification effectively achieve this, ensuring that the synthesized mRNA is suitable for downstream applications [16] [17].

Advancements in RNA purification techniques have significantly improved the quality of synthesized mRNA. Cellulose bead purification, in particular, has emerged as a reliable method for differentiating ssRNA from dsRNA contaminants. This process not only enhances RNA purity but also minimizes the risk of immune activation, which can compromise transfection efficiency and protein expression [18][19][20]. Furthermore, the integration of nanodrop spectrophotometry and gel electrophoresis enables accurate assessment of RNA yield and integrity. High-quality mRNA is characterized by distinct gel bands with minimal smearing, indicating optimal synthesis conditions. Notably, incubation times during IVT play a crucial role in determining RNA quality. Shorter incubation periods yield sharp RNA bands, while prolonged incubation often results in degradation and smearin [21].

The incorporation of pseudouridine (Ψ) into mRNA enhances its translational efficiency and stability, offering significant advantages over unmodified RNA. In this study, luciferase RNA synthesized with pseudouridine-modified rNTPs demonstrated superior stability and translational capacity compared to its unmodified counterpart. Functional validation experiments revealed a sevenfold increase in luciferase activity in cells transfected with pseudouridine-modified RNA, underscoring the importance of nucleotide modifications in therapeutic applications. Similar improvements were observed with GFP RNA, where pseudouridine-modified mRNA achieved higher fluorescence and protein expression levels. These findings align with previous research emphasizing the role of modified nucleotides in evading immune detection and enhancing RNA half-life [22] [23].

The cellular uptake and protein expression efficiency of mRNA are critical determinants of its therapeutic potential. HEK293 cells, a widely used model for transfection studies, were employed to evaluate the functional integrity of the synthesized mRNA. Transfection of GFP and luciferase mRNA into HEK293 cells revealed that cellulose-purified RNA exhibited superior transfection efficiency compared to conventionally purified RNA [24]. Fluorescence microscopy and luciferase activity assays confirmed that pseudouridine-modified mRNA consistently outperformed unmodified RNA in terms of protein expression. These results highlight the synergistic effects of nucleotide modifications and optimized purification protocols in achieving high-quality mRNA for therapeutic use [22].

Despite these advancements, challenges remain in the large-scale production and purification of therapeutic mRNA [10]. Double bands observed in gel electrophoresis suggest the presence of truncated RNA products or incomplete synthesis, which may impact the consistency and scalability of mRNA production [25][26]. Addressing these issues requires further optimization of reaction conditions and purification protocols. Additionally, the scalability of cellulose-based purification methods must be evaluated to ensure their feasibility for clinical-grade RNA production [22] [27] [28]. In conclusion, this study establishes a comprehensive framework for producing high-quality mRNA with enhanced stability and transfection efficiency. By integrating pseudouridine modifications, optimized IVT conditions, and advanced purification techniques, the synthesized mRNA achieves superior functional performance. These findings underscore the transformative potential of mRNA technology in therapeutic applications, from vaccines to protein-replacement therapies. Future research should focus on refining production protocols, scaling up manufacturing processes, and exploring novel delivery systems to maximize the clinical impact of mRNA-based therapeutics [29] [30] [31].

## MATERIALS AND METHOD

### 1. GFP /Luc DNA Template Preparation

The GFP DNA and Luc DNA templates were prepared separately and used for in vitro transcription (IVT). The GFP DNA construct (T7 promoter, kozak seqence, GFP coding sequence, his-tag and polyA tail) was used as template for GFP IVT. In the same way the Luc DNA construct (T7 promoter, kozak seqence, Luciferase coding sequence and polyA tail) was used as template for Luciferase IVT. Both the templates were amplified using PCR. The PCR amplified DNA was subjected to run gel electrophoresis cut the desired band; use the gel purification kit to get highly purified DNA prior to IVT.

### 2. In Vitro Transcription (IVT) Reaction

IVT was performed using the GFP DNA template under different incubation times (3 hours, 7 hours, and 16 hours). The reaction included rNTPs and pseudouridine triphosphate (ΨTP) as nucleosides. For the generation of nucleoside-modified mRNAs, uridine 5’-triphosphate (UTP) was replaced with the pseudouridine triphosphate (ΨTP) in the transcription reaction. Capping of the IVT mRNAs was performed co-transcriptionally using the ARCA cap. After DNase digestion, the synthesized mRNA was purified. The IVT reaction was carried out in RNase-free conditions using the following components in total 20 μL reaction mixture:

- DNA template: 2 μL (200 ng)
- 10X transcription buffer: 2 μL
- 100 mM ATP: 2 μL
- 100 mM UTP or pseudouridine (Ψ): 2 μL
- 100 mM CTP: 2 μL
- 20 mM GTP: 2 μL
- 100 mM anti-reverse cap analog (ARCA): 0.8 μL
- 100 mM DTT: 2 μL
- RNase inhibitor: 2 μL
- T7 RNA polymerase: 2 μL
- RNase-free water: 1.2 μL

### 3. Purification of Synthesized mRNA

The synthesized mRNA was purified to remove dsRNA contaminants, which can activate TLR3 receptors, and residual DNA templates. DNase digestion was performed before column-based purification. Cellulose bead purification was used to separate ssRNA from dsRNA. The quality and yield of purified RNA were assessed using Nanodrop spectrophotometry and gel electrophoresis. The RNA concentration was determined using a Nanodrop spectrophotometer. Aliquots of IVT mRNAs were analyzed by electrophoresis in 1% agarose gels containing 0.5μg/mL EtBr for nucleic acid gel stain.

RNA purification was performed as follows:

- Luc-RNA was synthesized by IVT and treated with DNase,
- The total RNA was treated with Cellulose bead in HEPE+ 15% ETOH,
- Spine down the cellulose, ss-RNA was eluted from supernatant and ds-RNA from cellulose,
- The eluted supernatant containing ss-RNA was purified using spin column RNA purification kit.
- RNAs were checked in nanodrop, and gel electrophoresis. Concentration measurements showed minimal cellulose-bound RNA, while total RNA and ssRNA displayed consistent levels. RNA samples were analyzed via gel electrophoresis.

### 4. Modified and Unmodified RNA Preparation

Luciferase RNA was transcribed with both normal rNTPs and pseudouridine (Ψ)-modified+ rNTPs separately using DNA template (T7 promoter + kozak + Luc + polyA). The reaction mix was performed according to the above protocol with 3hrs incubation. DNase digestion was performed one/twice to eliminate DNA contaminants. Purification was completed using cellulose bead based methods to distinguish ssRNA and potential dsRNA contaminants.

RNA purification was performed as follows:

- RNA was mixed with Cellulose bead in HEPE+ 15% ETOH.
- Centrifuged the mixture.
- Isolate supernatant followed by spin column purification of ss-RNA.

The quality and yield of purified RNA were assessed using Nanodrop spectrophotometry and gel electrophoresis.

### 5. IVT mRNA Transfection in cells

500ng of purified mRNA encoding GFP / luciferase (both modified and unmodified) was transfected into HEK293 cells using Lipofectamine™ RNAiMAX Transfection Reagent in 96-well plates according to manufacturer recommendation. The cells were incubated in CO_2_ incubator for 24 hrs.

### 6. Protein Analysis

For Luciferase activity, after 24 hrs of incubation the growth medium was removed from cultured cells, washed cells in 1X PBS, dispensed a 20μl/well of 1X lysis reagent for a 96 well plate into. Pellet debris by brief centrifugation, and transfer the supernatant to a new tube. Mix 20μl of cell lysate with 100μl of Luciferase Assay Reagent and the light was measured in Plate-Reading Luminometer.

After incubation GFP expression was assessed using fluorescence microscopy and Western blot analysis with His-tag antibodies. Shortly, after 24 hrs of cell culture, cell lysate were made in RIPA buffer. Samples were run on a 10% SDS PAGE, and Western blotting was performed using standard techniques with anti His-antibody.

## RESULTS

### 1. GFP RNA Synthesis Efficiency

In vitro transcription (IVT) of GFP RNA was analysed at 3, 7, and 16-hour incubation intervals. Gel electrophoresis analysis demonstrated that a 3-hour incubation period yielded sharp RNA bands with minimal smearing, indicating optimal synthesis conditions. Longer incubation times (7 and 16 hours) resulted in reduced RNA quality, characterized by increased smearing (Figure 1,B). Consequently, a 3-hour incubation period was selected for subsequent experiments.

**Figure 1:**
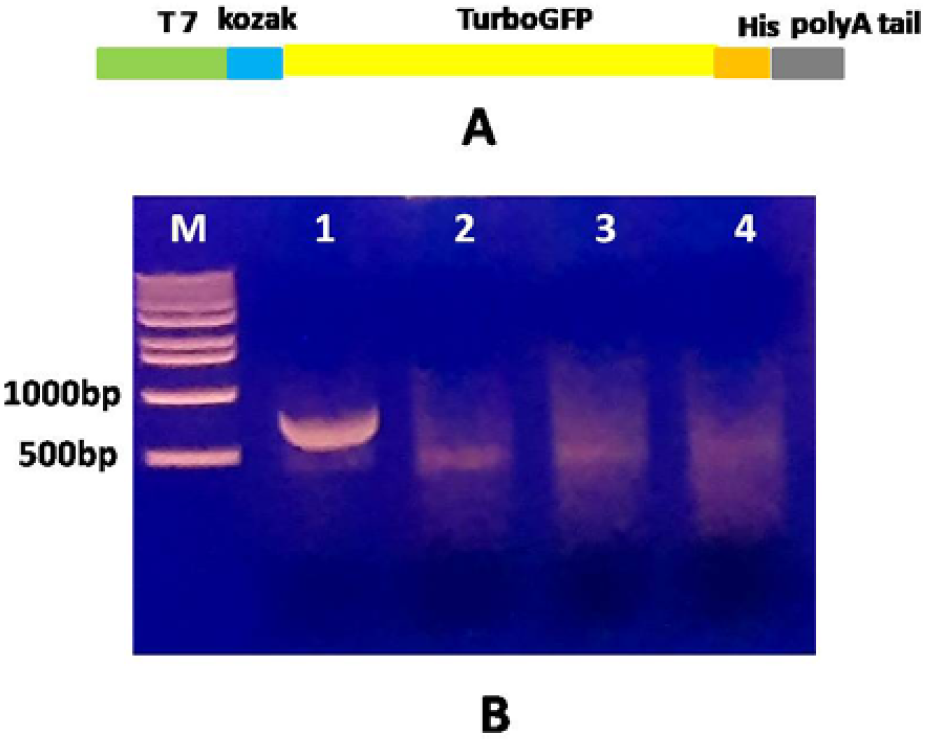
(A) Used template for IVT (B) Gel electrophoresis of IVT mRNA (GFP) in different incubation time. GFP DNA template (Lane-1), GFP RNA by IVT for 3 hrs (Lane-2), GFP RNA by IVT for 7 hrs (Lane-3), GFP RNA by IVT for 16 hrs (Lane-4).

### 2. RNA Purification Efficacy

The purification of in vitro transcribed (IVT) mRNA requires effective removal of contaminants such as reaction buffers, template DNA, and double-stranded RNA (dsRNA) to ensure quality and functionality. During IVT, dsRNA is co-synthesized with single-stranded mRNA (ssRNA) and must be eliminated because dsRNA activates TLR3 receptors, triggering unwanted immune responses. Template DNA is typically degraded using DNase before column-based purification, while dsRNA removal is achieved through cellulose-based treatments, which selectively bind dsRNA but not ssRNA. Studies comparing purification methods show that cellulose treatment effectively separates ssRNA (retaining high purity) from dsRNA, with minimal dsRNA detected in purified samples. This optimized protocol significantly reduces dsRNA contamination, producing high-quality mRNA suitable for downstream applications like therapeutic development or research.

To standardize RNA purification, we used a PCR-amplified DNA template containing a T7 promoter, Kozak sequence, luciferase coding region, and a polyA tail (Figure 2). RNA was synthesized using either ARCA cap with unmodified rNTPs or ARCA cap with modified rNTPs. Following DNase digestion, the synthesized RNA underwent cellulose bead treatment, and was then further purified using column purification.

**Fig 2:**
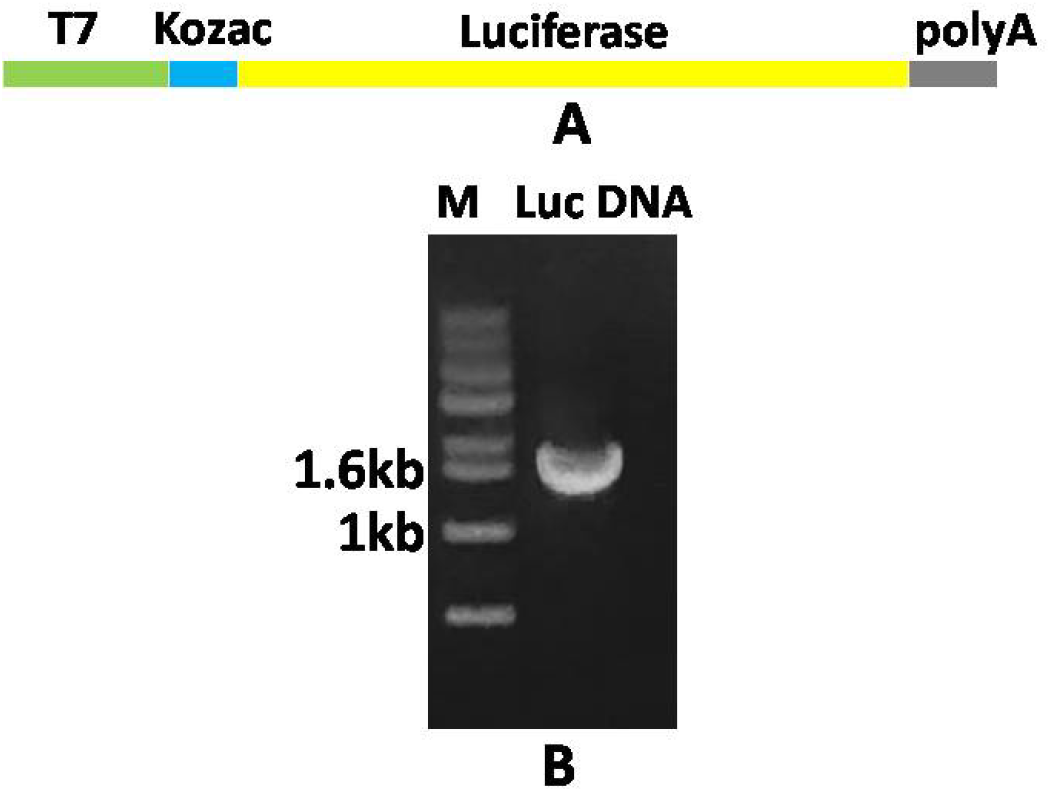
PCR amplified Luc DNA template containing T7 promoter, Kozac sequence, Luciferase coding sequence and polyA tail (A). PCR amplified Luc DNA template containing T7 promoter, RBS, Luciferase coding sequence and polyA tail (B).

All RNA samples were analyzed by gel electrophoresis. Cellulose-bound RNA appeared in lanes 3 and 4 (Figure 3), indicating the presence of dsRNA. In contrast, column-purified RNAs displayed double bands. To assess the functional efficacy of these RNAs, we transfected cells with the purified samples and measured luciferase activity. Notably, RNA containing Ψ modification showed a seven-fold increase in luciferase activity compared to unmodified control RNA (Figure 4).

**Fig 3:**
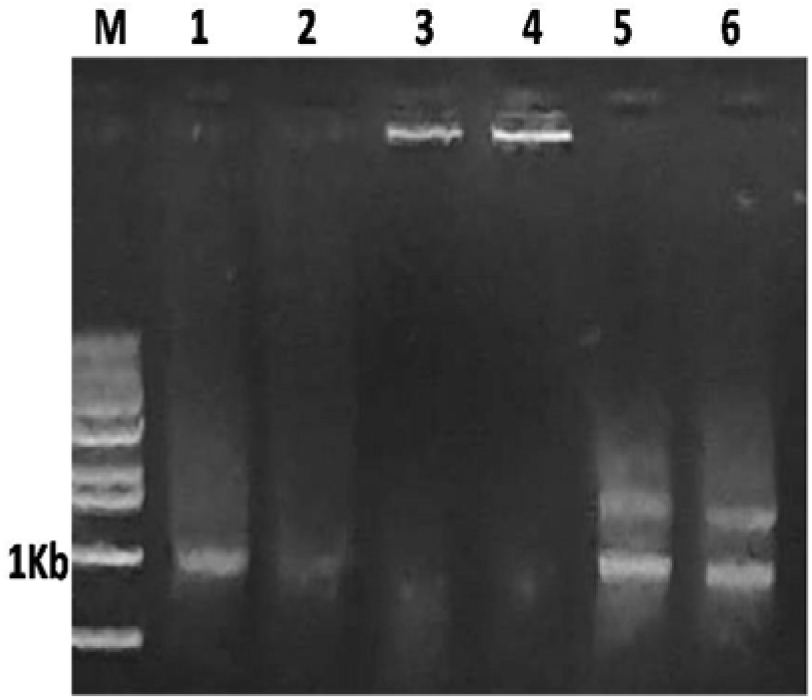
Cellulose-based purification of luciferase mRNA, 1. Luc transcript with ARCA +unmodified rNTPs after DNase digestion, 2. Luc transcript with ARCA+rATP+rGTP+Ψ+rCTP after DNase digestion, 3. Cellulose bound RNA of lane1 (might be dsRNA, showing in the well), 4. Cellulose bound RNA of lane2 (might be dsRNA, showing in the well), 5. Cellulose based Purified RNA of lane 1, 6. Cellulose based Purified RNA of lane 2.

**Fig 4:**
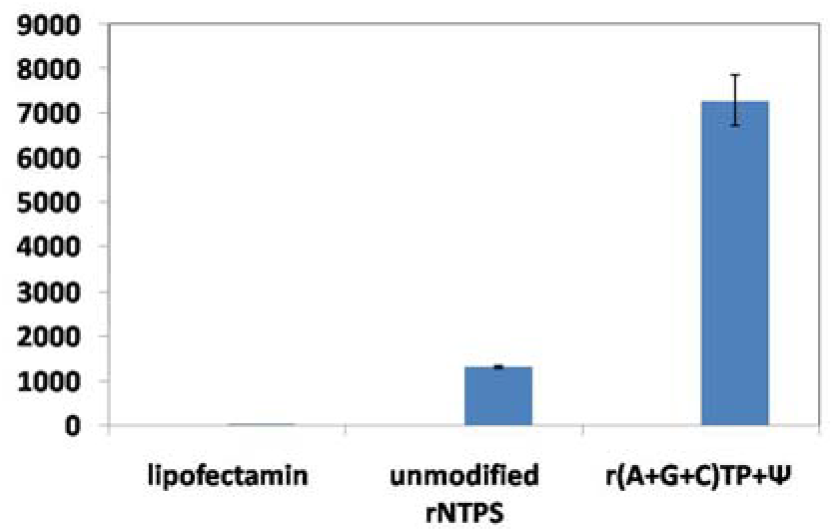
Luciferase activity of unmodified and modified RNA, Transfection was done with cellulose bead purified RNA in HEK293 cells. The result is showing 7 times increased luciferase activity was found in Ψ compared to control (unmodified RNA).

To investigate the double bands observed in the purified RNAs, we performed an additional DNase digestion. However, this treatment did not alter the banding pattern in gel electrophoresis (Figure 5).

**Fig 5:**
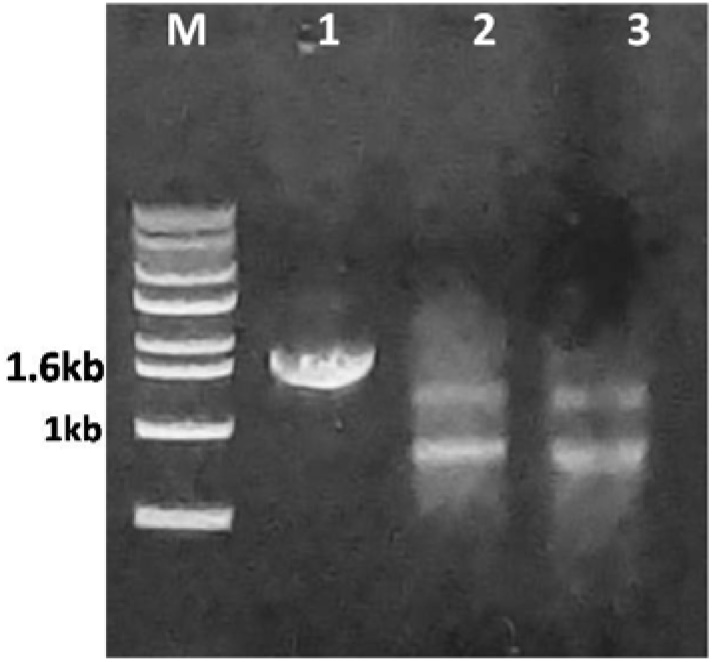
Gel electrophoresis showing Luciferase RNA with 2 times DNase digestion, 1. PCR amplified Luc DNA, 2. Luc transcript with unmodified rNTPs after2nd DNase treatment, 3. Luc transcript with rATP+rGTP+Ψ+rCTP after 2nd DNase treatment.

#### Modified vs. Unmodified RNA Analysis

Luciferase RNA synthesized with pseudouridine (Ψ)-modified rNTPs exhibited improved stability compared to unmodified RNA. Gel electrophoresis revealed distinct bands for dsRNA after cellulose-based purification, while modified RNA showed enhanced integrity post-DNase digestion. The stability of modified RNA supports its suitability for therapeutic and research applications.

However, luciferase activity assays confirmed the functional integrity of the synthesized mRNA. Further analysis is required to determine whether the secondary band represents truncated RNA products or incomplete synthesis. Purified mRNA with pseudouridine modifications exhibited significantly higher luciferase activity compared to unmodified mRNA, underscoring the importance of RNA modifications in enhancing translation efficiency.

**Fig 6:**
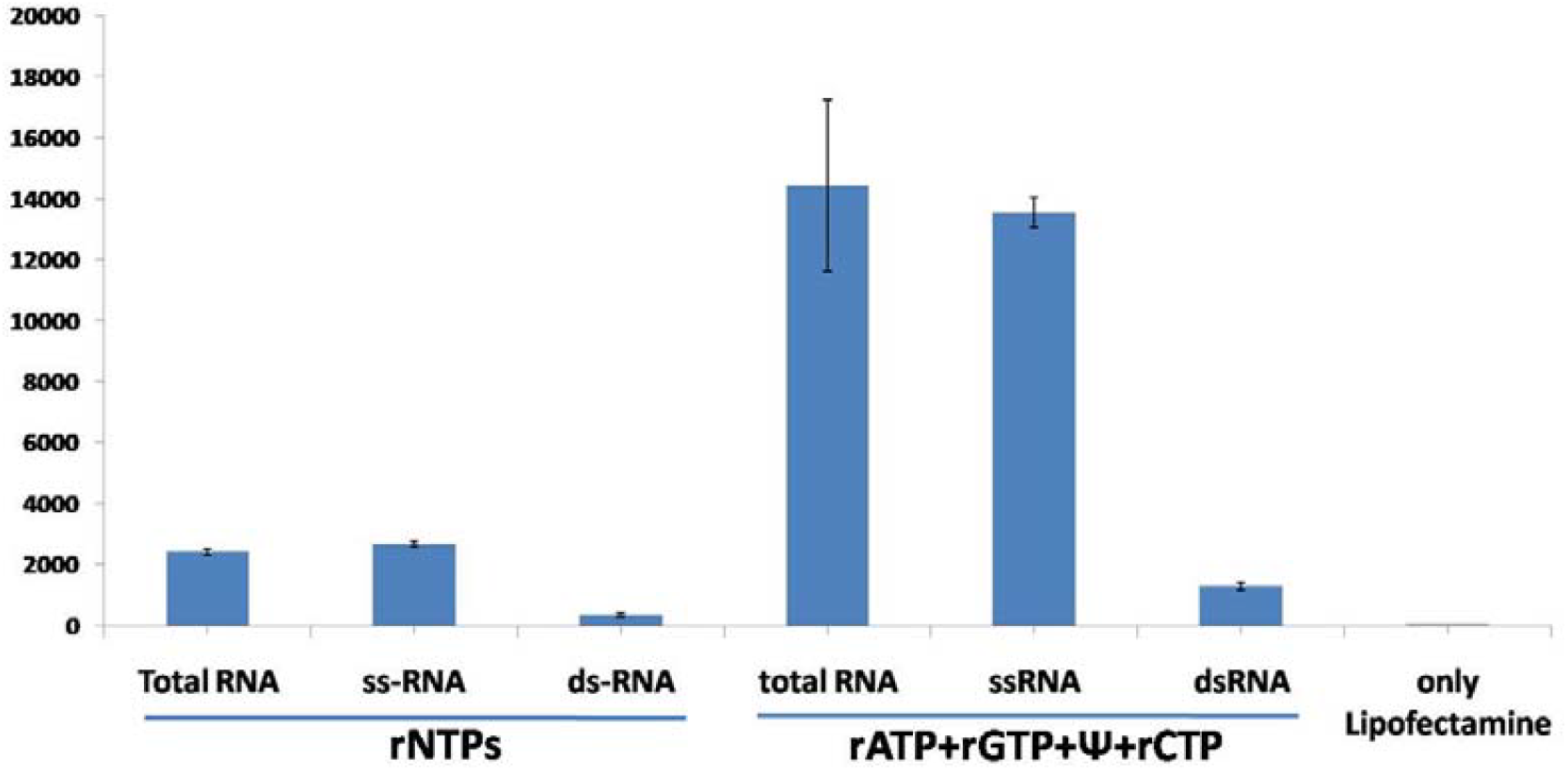
Luciferase assays of isolated total RNA, ss-RNA and ds-RNA.

### 3. GFP mRNA Transfection and Expression

Transfection experiments using GFP RNA in HEK293 cells highlighted the superior efficiency of cellulose-purified RNA. Modified GFP RNA showed higher fluorescence compared to unmodified RNA, confirming enhanced expression. Similarly, luciferase activity assays revealed a 7-fold increase in pseudouridine-modified RNA transfections, indicating improved translation efficiency and cellular uptake.

GFP protein expression (26.76 kDa) was validated via fluorescence imaging and western blot. Transfections with modified RNA produced significantly higher GFP protein levels, aligning with the observed fluorescence. The presence of His tags facilitated protein purification and ensured accurate downstream analyses.

**Fig 7:**
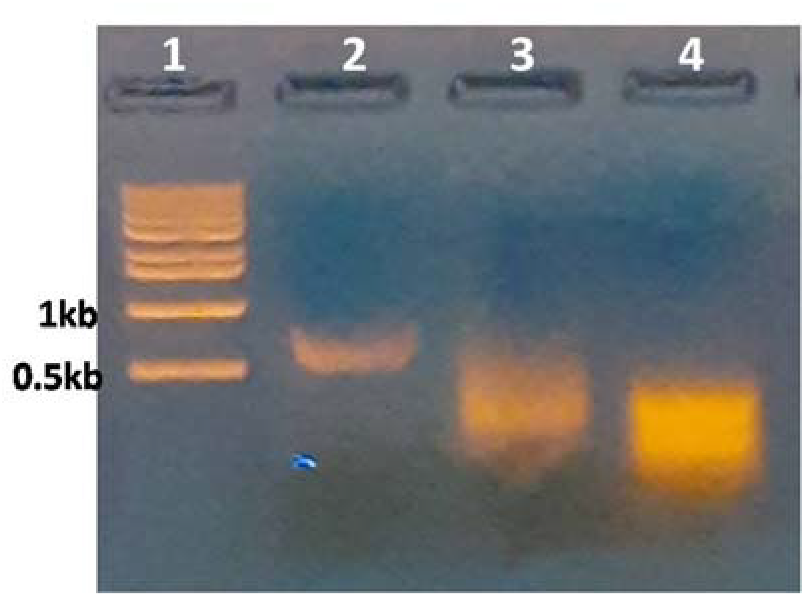
RNA gel electrophoresis of IVT mRNA 1: M, 2-DNA template, 3: IVT mRNA with ARCA+rNTPs, & 4: ARCA+ rATP+rGTP+Ψ+rCTP mRNA.

**Fig 8:**
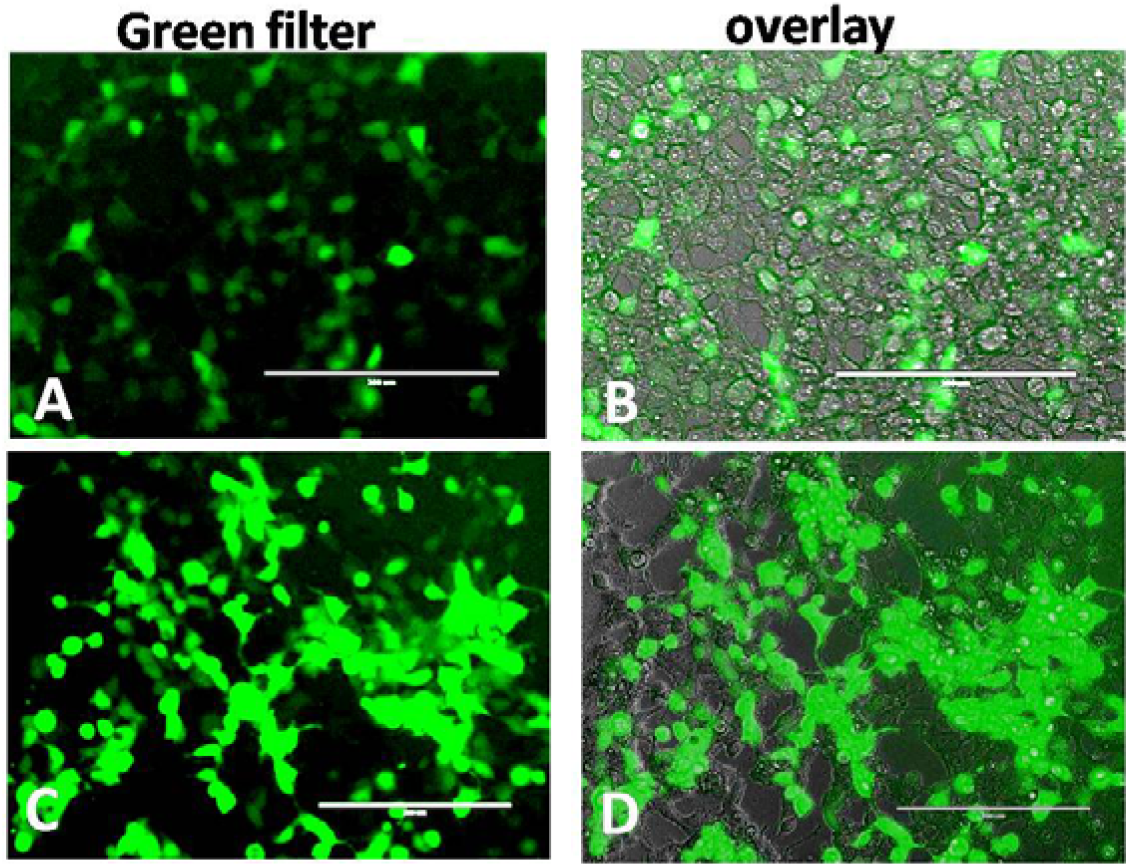
Expression of GFP in HEK293 cells, a&b: rNTPs GFP mRNA transfection, c&d: r(A+C+ Ψ+G)TPs GFP mRNA transfection.

**Fig 10:**
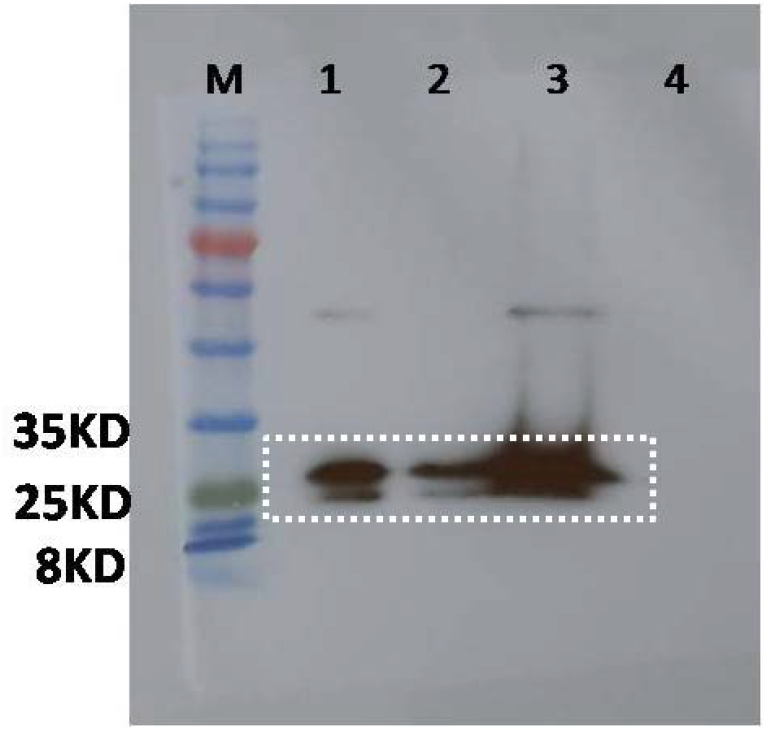
Western Blotting of isolated protein from GFP mRNA transfected cells (HEK293) using His tag antibody (genescript), 1&2: rNTPs protein samples 3: r(A+G+C)TPs+Ψ protein sample 4: Control cell protein (without transfection).GFP contain His tag at C-terminal end showing protein band with His antibody, GFP (26.7kd)

#### Enhanced Luciferase Expression by Pseudouridine and Its Combinations Indicates Improved Translational Efficiency

Chemically modified luci mRNA were synthesized using various nucleoside analogs and assessed for their transfection efficiency based on luciferase activity. Among all the modifications tested, pseudouridine (Ψ) substitution exhibited the highest luciferase activity, significantly surpassing the native unmodified mRNA, suggesting enhanced translation efficiency and mRNA stability.

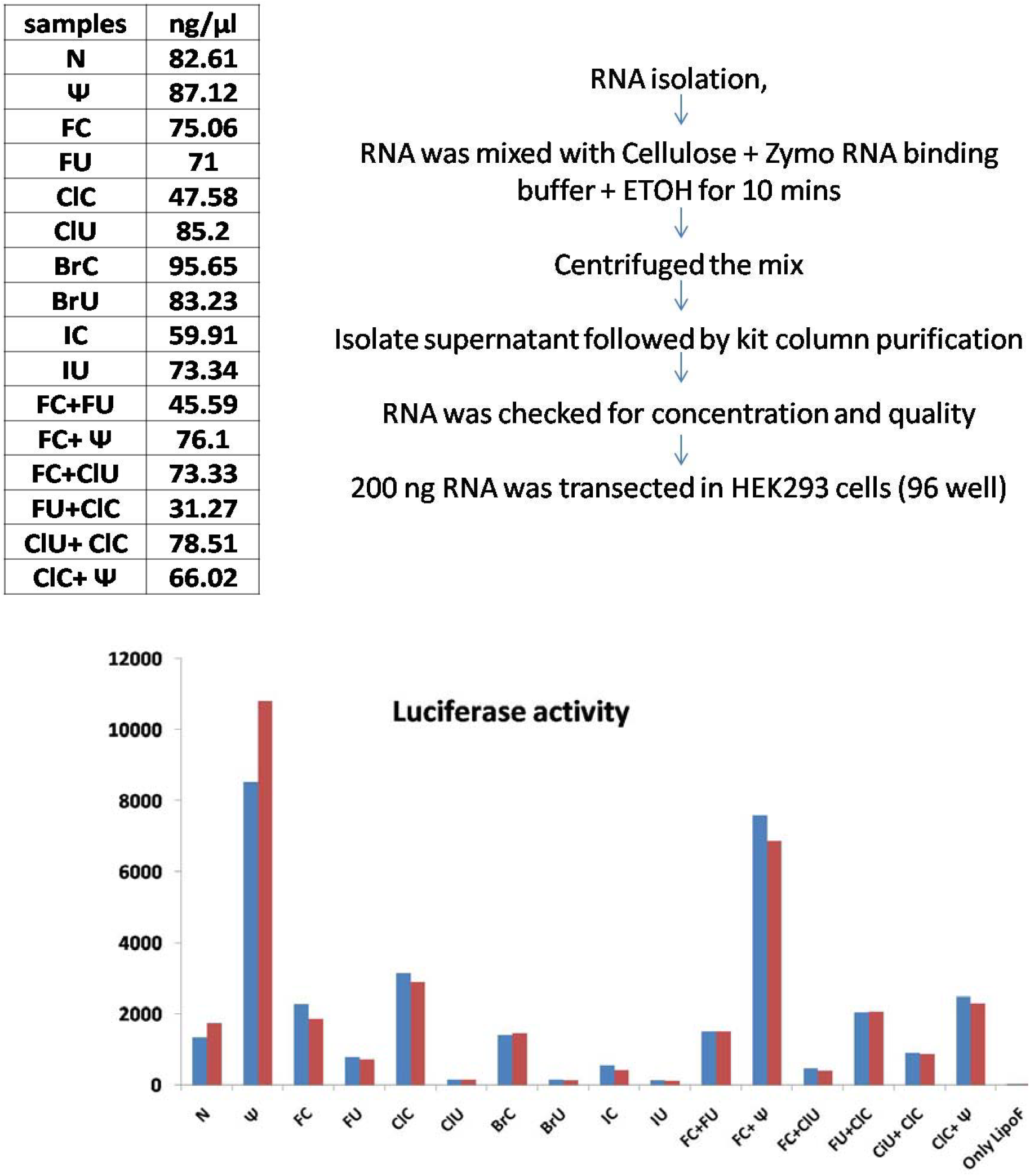

Samples names-Luciferase mRNA containing all Natural: N, pseudorUTP: Ψ, fluoro-rCTP: FC, 5-fluoro-rUTP: FU, 5-chloro-rCTP: CLC, 5-chloro-rUTP: CLU, 5-bromo-rCTP: BrC, 5-bromo-rUTP: BrU, 5-iodo-rCTP: IC, 5-Iodo-rUTP: IU, Pseudo-rUTP and 5-fluoro-rCTP: Ψ +FC, pseudo-rUTP and 5-chloro-rCTP: Ψ+CLC, 5-fluoro-rCTP and 5-chloro-rUTP: FC+CLU, 5-fluoro-rUTP and 5-chloro-rCTP: FU+CLC, 5-fluoro-rCTP and 5-fluoro-rUTP: FC+FU, 5-chloro-rCTP and 5-chloro-rUTP: CLC+CLU.

## DISSICUSSION

The findings of this study highlight the transformative potential of pseudouridine (Ψ)-modified RNA in improving RNA-based applications. The enhanced stability of modified RNA, as evidenced by distinct gel electrophoresis patterns and reduced degradation, underscores its potential for therapeutic use. This aligns with previous research emphasizing the role of nucleotide modifications in evading immune detection and prolonging RNA half-life [14]. Cellulose-based purification emerged as a critical step in removing dsRNA contaminants, a known cause of innate immune activation and reduced transfection efficacy. The methodology provided a high yield of ssRNA, suitable for effective gene expression, as confirmed by the significant increase in GFP fluorescence and luciferase activity in HEK293 cells. Such improvements demonstrate the method’s applicability for clinical-grade RNA production.

The 7-fold increase in luciferase activity with Ψ-modified RNA highlights the importance of nucleotide modifications in enhancing translational efficiency [32]. These results support its potential for mRNA vaccine development, where robust protein expression is crucial for eliciting an immune response. However, challenges remain, including gel electrophoresis anomalies, such as double bands, which suggest incomplete RNA processing or co-purification of RNA species. Further investigations are needed to optimize reaction conditions and improve RNA integrity. Addressing these issues will ensure the reproducibility and scalability of this approach for broader applications.

Luciferase assay results revealed a significantly higher expression level in cells treated with the pseudouridine (Ψ) sample, showing maximal activity compared to other analogs. The combination of FC and Ψ also resulted in enhanced luciferase expression, suggesting a possible synergistic effect.

In conclusion, this study establishes a framework for producing high-quality, modified RNA with enhanced stability and transfection efficiency. Future work will focus on scaling up the process, improving purification protocols, and exploring its use in diverse therapeutic contexts, including vaccines and RNA-based therapeutics.

## Author contribution

Subrata Kumar Das: Conceptualization, Supervision, Methodology, Investigation, Formal analysis, Data curation, Visualization, Validation, writing, review & editing,. Gaurav Dutt: Investigation, writing, review & editing, Vishakha Goswami: review & editing Abul Faiz: review & editing and Alpana Joshi: review & editing.

## Acknowledgments

This research was supported by the Professor Young Jun Seo’s laboratory, Department of Chemistry, Jeonbuk National University, Jeonju 54896, South Korea.

## Declaration of competing interest

The authors declare that they have no known competing financial interests or personal relationships that could have appeared to influence the work reported in this paper.

## Notes

### Competing Interest Statement

The authors have declared no competing interest.

